# Live-Attenuated CHIKV Vaccine with Rearranged Genome Replicates *in vitro* and Induces Immune Response in Mice

**DOI:** 10.1101/2023.09.16.558061

**Authors:** Irina Tretyakova, Joongho Joh, Igor S. Lukashevich, Brian Alejandro, Mary Gearon, Donghoon Chung, Peter Pushko

## Abstract

Chikungunya fever virus (CHIKV) is a mosquito-borne alphavirus that causes wide-spread human infections and epidemics in Asia, Africa and recently, in the Americas. There is no approved vaccine and CHIKV is considered a priority pathogen by CEPI and WHO. Previously, we developed immunization DNA (iDNA) plasmid capable of launching live-attenuated CHIKV vaccine *in vivo*. Here we report the use of CHIKV iDNA plasmid to prepare a novel, live-attenuated CHIKV vaccine V5040 with rearranged RNA genome for improved safety. In V5040, genomic RNA was rearranged to encode capsid gene downstream from the glycoprotein genes. To secure safety profile, attenuated mutations derived from experimental CHIKV 181/25 vaccine were also engineered into E2 gene of V5040. The DNA copy of rearranged CHIKV genomic RNA with attenuated mutations was cloned into iDNA plasmid pMG5040 downstream from the CMV promoter. After transfection in vitro, pMG5040 launched replication of V5040 virus with rearranged genome and attenuating E2 mutations. Furthermore, V5040 virus was evaluated in experimental murine models for safety and immunogenicity. Vaccination with V5040 virus subcutaneously resulted in elicitation of CHIKV-specific, virus-neutralizing antibodies. The results warrant further evaluation of V5040 virus with rearranged genome as a novel live-attenuated vaccine for CHIKV.

## INTRODUCTION

Chikungunya virus (CHIKV) is a mosquito-borne alphavirus that causes epidemics of chikungunya fever. Outbreaks of CHIKV are continuing worldwide [1]. CHIKV is transmitted to humans primarily by *Aedes aegypti* mosquito [2, 3]. In addition, during the 2005–2006 epidemic in Reunion islands in the Indian Ocean that affected more than one-third of the island population and caused 284 deaths, a new mosquito vector *A. albopictus*, was identified [3, 4]. Outbreaks of CHIKV included India in 2005–2006 with estimated 1.3 million people infected [5, 6] and the Americas since 2014, with more than 150,000 confirmed CHIKV infections in the Caribbean region only [7]. CHIKV is also widespread in Africa and South East Asia [7, 8]. With an increase in global travel, the risk for rapid expansion of CHIKV to non-endemic areas has increased [9, 10]. Some travelers are viremic, and *A. albopictus* is common in urban areas of the U.S., raising concerns for the immunologically naïve population [11]. Local transmission cases have been reported in Europe and the U.S. [11-15]. Changing climate patterns also favor geographical expansion of CHIKV [16]. Given the current large outbreaks and the worldwide distribution of *A. aegypti* and *A. albopictus*, CHIKV represents a global public health threat [1].

CHIKV causes fever, headache, rash, nausea, myalgia, and arthralgia [17]. Complications include respiratory failure, cardiovascular disease, hepatitis, cutaneous effects, and central nervous system problems [18, 19]. More than 50% of patients who suffer from severe infection are over 65 years old, and more than 33% of them die. Therefore, CHIKV vaccine is needed. Currently, there is no approved CHIKV vaccine or specific antiviral therapy in the U.S. Available therapeutic interventions include anti-inflammatory drugs, fluids, and bed rest. Antivirals and agents that restrict the cell-to-cell spread of the virus can be useful, but not FDA-approved [20]. Arthralgia associated with the fever can persist for months or years and progresses to arthritis in some patients [21].

Although no CHIKV vaccine has been approved, several live-attenuated vaccines have been evaluated in the late clinical trials [22]. These included vaccines with attenuating mutations in the nonstructural and structural genes. An investigational CHIKV vaccine clone 181/25 has been immunogenic in a Phase II clinical trial; however, mild transient arthralgia was observed in some vaccinated individuals, indicating the need for safety improvement [23]. Addressing the need for a safe and efficacious vaccine, we previously developed a novel plasmid DNA that expressed the full-length genomic RNA of live-attenuated CHIKV from a eukaryotic promoter - a strategy termed iDNA [24]. The iDNA plasmid can be transfected in cell culture to manufacture live-attenuated virus vaccine, or can be administered directly into patients’ tissues to generate live-attenuated virus *in vivo*. In the latter case, iDNA plasmid is taken up by a limited number of cells, and CHIKV genomic RNA is transcribed in these cells. As a result, CHIKV proteins are synthesized, genomic RNA is packaged into virus particle, and live-attenuated CHIKV is assembled and secreted from the cells. In immunocompetent BALB/c mice, the prototype 181/25-based iDNA plasmid vaccine demonstrated safety and protective efficacy [25].

In attempts to further improve CHIKV vaccine, here we have used CHIKV iDNA plasmid as a reverse genetics system to engineer novel CHIKV vaccine. The V5040 CHIKV with a rearranged RNA genome was prepared to improve safety. As it was documented for numerous RNA viruses, rearrangement of viral genes result in attenuation and resistance to reversions to wild type phenotypes [26]. We placed CHIKV capsid gene downstream the glycoprotein gene under control of a duplicate subgenomic 26S promoter. In addition, two attenuating mutations derived from the prototype 181/25 vaccine were included in the E2 glycoprotein region. We report that live virus with rearranged genome replicates after pMG5040 plasmid is transfected in Vero cells. Thus, rearranged CHIKV iDNA pMG5040 launches live-attenuated CHIKV V5040 in Vero cell culture. Furthermore, vaccination of mice with V5040 confirmed its safety and elicited CHIKV-specific, virus-neutralizing serum antibodies. We conclude that V5040 vaccine is a promising vaccine for CHIKV, and further evaluation is warranted.

## MATERIALS AND METHODS

### Cells and Plasmids

Vero cell line (American Type Culture Collection, Manassas, VA) was maintained in a humidified incubator at 37°C in 5% CO_2_ in αMEM supplemented with 10% fetal bovine serum (FBS) and gentamicin sulfate (10 μg/ml) (Life Technologies, Carlsbad, CA).

The pMG5040 plasmid was derived from iDNA plasmid p181/25 described elsewhere [25]. The pMG5040 was based on pUC backbone vector and encoded the full-length rearranged RNA genome of V5040 vaccine virus (Figure 1). The capsid (C) gene was cloned downstream from the glycoprotein gene and expressed using the duplicate subgenomic promoter. The ATG codon was introduced at the 5’ of E2 gene within the glycoprotein (GP) genes, while a TGA stop codon was introduced at the 3’ of the C gene. The full-length genomic cDNA of V5040 virus was placed in the pMG5040 under transcriptional control of the optimized CMV promoter. The hepatitis delta ribozyme was introduced downstream from the V5040 cDNA to ensure cleavage of the genomic RNA transcript after the synthetic poly(A) at the viral 3’ end. The cDNA of V5040 also maintained attenuating mutations Thr12Ile and Gly82Arg (both in the E2 gene) derived from the 181/25 prototype vaccine sequence.

**Figure 1.**
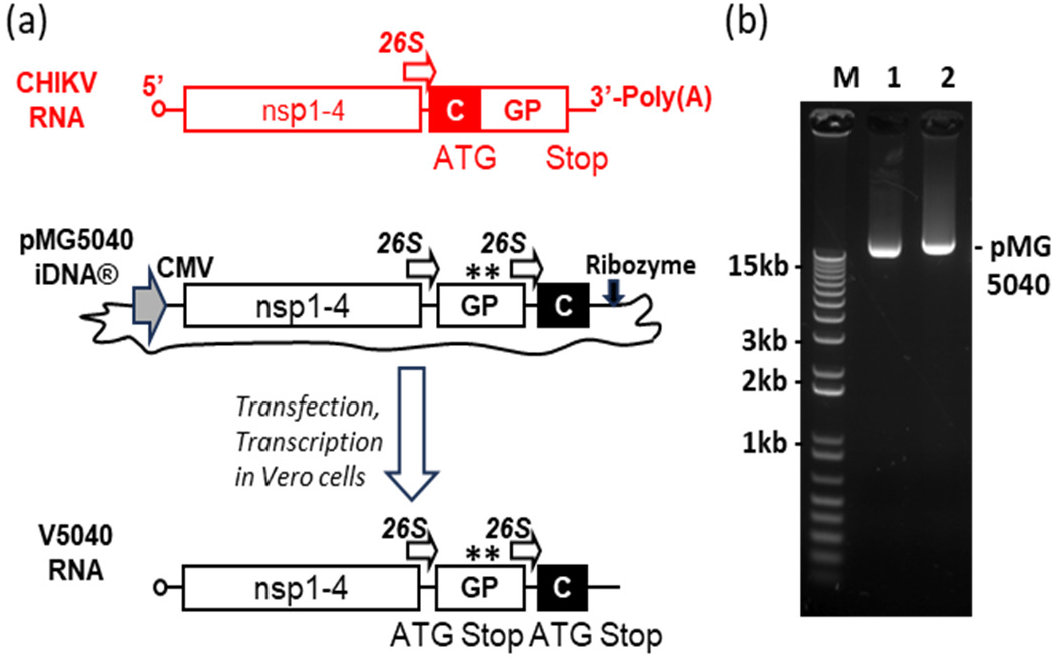
Genetic maps depicting the pMG5040 plasmid encoding the full-length rearranged RNA. (a) Indicated are approximate locations of cytomegalovirus (CMV) promoter (solid arrow), 26S promoter (open arrows) and attenuating mutations (asterisks). The wild type CHIKV genome is shown in red. CHIKV pMG5040 plasmid encoded rearranged full-length infectious CHIKV RNA under transcriptional control of the CMV promoter. The plasmid pMG5040 contained both attenuating mutations (asterisks) from the IND vaccine 181/25. (b) 1% Agarose/TAE gel showing pMG5040 plasmid as compared to control VEEV vaccine. M, 1kB Plus DNA ladder (Thermo); 1, VEE control pMG4020: 2, CHIKV pMG5040.

The iDNA plasmid pMG5040 containing the rearranged, full-length CHIKV iDNA was isolated from *E. coli*. The CHIKV sequence was confirmed by DNA sequencing. Bioinformatic analysis was performed to ensure that rearranged CHIKV sequences do not interfere with plasmid production in *E. coli* or the translation of the viral genome in eukaryotic cells. We confirmed the presence of the authentic ORFs and the absence of any strong internal bacterial promoter and transcription sites that can lead to expression of potentially toxic proteins and inhibit plasmid growth in *E coli*. The ORFs were predicted by using NCBI ORF Finder software [27]. Potential bacterial promoters were predicted by using BPROM software [28], which is a bacterial promoter recognition program with approximately 80% accuracy and specificity. In addition, rearranged sequence was screened for potential splice sites that can lead to degradation of RNA in the nucleus, using software developed within Berkeley Drosophila Genome Project [29]. The amino acid sequence of CHIKV proteins was kept according to GenBank 181/25 TSI-GSD-218 CHIKV 181/25 vaccine #L37661 except additional Met at the start of GP genes, and a stop codon at the end of C gene (Figure 1).

The resulting plasmid pMG5040 was propagated in *E. coli* Stbl3 cells (Thermo, Carlsbad, CA) using standard Luria Broth LB medium in the presence of kanamycin. Plasmid was isolated by an endotoxin-free DNA isolation method (Qiagen, Valencia, CA), or a similar DNA isolation method, according to manufacturer’s instructions. Finally, pMG5040 was formulated in phosphate-buffered saline (PBS) to a concentration of ∼1 mg/ml. This process resulted in a transfection-grade, sterile DNA with 95% supercoiled DNA and an A_260_/A_280_ ratio of ∼1.9, as well as minimal residual endotoxin, RNA, genomic DNA, and protein impurities.

### Viruses

Live-attenuated vaccine virus V5040 was prepared by transfecting Vero cells with pMG5040 plasmid. Transfection was done by electroporation essentially as described previously [25, 30]. Briefly, Vero cells were transfected with 100 ng of pMG5040 and incubated at 37°C. The transfection medium was harvested at 48 h post-transfection (h.p.i.) and filter-sterilized using 0.22 uM filter. The virus was then concentrated and partially purified by ultracentrifugation and resuspended in phosphate buffered saline (PBS), pH7.4. The V5040 titer was determined by a standard plaque assay in Vero cell monolayers in 6-well tissue culture plates using serial dilutions of the virus, and stained using neutral red. To detect the virus in samples with low titers (below the limit of direct plaque assay, 25 PFU/ml) co-cultivation of test samples with Vero cells was used to amplify the low viral load. The transfection medium was used to infect Vero cells at a multiplicity of infection of 0.01 to generate V5040 passage 1 (P1) virus. The P1 virus was harvested at 48 h.p.i. and the titer of the V5040 P1 virus was determined. The virus was aliquoted and stored at -80°C until used *in vitro* or *in vivo*.

To determine virus growth kinetics, Vero cells were infected in 75 cm^2^ flasks with 100 PFU or 1000 PFU of V5040 virus, or CHIKV 181/25 control virus, as described in the Results. In the same experiment, Vero cells were also transfected with 100 ng of pMG5040 by electroporation. Samples of the growth medium were taken at 12 h intervals.

As described above, CHIKV 181/25 vaccine virus was used as a control. The CHIKV 181/25 vaccine was received from the World Reference Center for Emerging Viruses and Arboviruses (WRCEVA) through the University of Texas Medical Branch (UTMB) in Galveston, Texas.

To detect neutralizing antibody, molecular clone-derived CHIKV 181/25 clone expressing nLuc (CHIKV 181/25-nLuc) was generated by transfecting infectious RNA into cells. Briefly, the nLuc gene was inserted within the nsP3 gene by Gibson cloning method at 5217 b.p. site. The backbone and the insert were obtained by PCR with pTD181_25, a plasmid with the full genome of CHIKV 181/25 provided from Dr. Kevin Sokoloski as a gift, and with a plasmid containing nLuc gene. Primer sets were used with Fusion high fidelity DNA polymerase (NEB M0530L) with the manufacturer-recommended protocol and standard molecular cloning techniques (Table 1). Constructed clones were validated by a whole plasmid sequencing were used as a template to generate infectious RNA by using in vitro transcription (ThermoFisher SP6 mMessage kit) after a linearization with Not I enzyme. Infectious virus was rescued from Vero76 cells transfected with the infectious in vitro transcribed RNA using Lipofectamine mMessenger max. Stock virus was generated by amplyfing the rescued virus once in Vero76 and stored at -80 °C in aliquots until used.

**Table 1.**
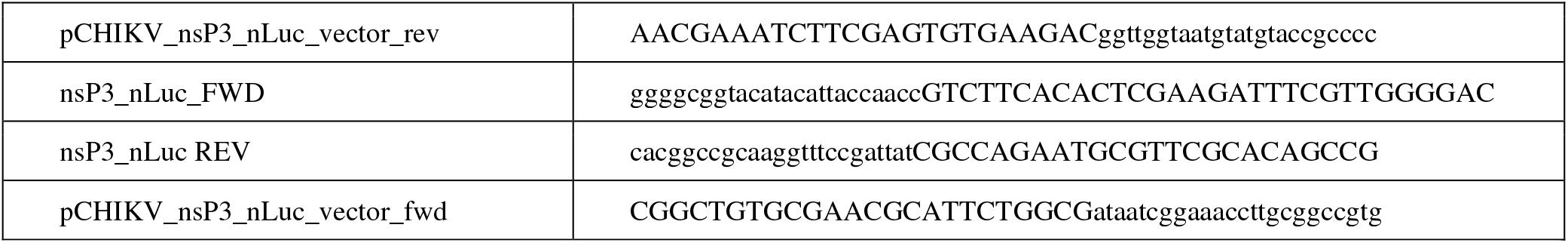
Primers used for preparation of CHIKV 181/25 clone expressing nLuc (CHIKV 181/25-nLuc). The nLuc gene was inserted within the nsP3 gene by Gibson cloning method at 5217 bp site as described in Materials and Methods.

Challenge was performed by inoculating a wild type CHIKV strain SL15649. Strain SL15649 was originally isolated from a CHIKV-infected patient in Sri Lanka in 2006, has been minimally passaged in cell culture, and is pathogenic in a mouse model of CHIKV disease [31].

### Immunizations of BALB/c mice

Research involving mice was done according to approved institutional animal protocols according to American Veterinary Medical Association recommendations. BALB/c mice (4-8 week-old, Noble Life Sciences, Woodbine, MD) were anesthetized with isoflurane prior to vaccinations. Mice (n = 5 to 8 animals per group, female, two independent repeats) were vaccinated subcutaneously (s.c.) with V5040 vaccine virus, or with control 181/25 vaccine in the dorsal area at the doses 10^4^ and 10^5^ PFU as indicated in the Results section. After vaccinations, animals were observed daily for signs of infection, morbidity and discomfort. Blood samples were collected from the retro-orbital sinus on days 0 (pre-bleed), day 2-4, and day 28. Viremia was evaluated on days 2-4 post-vaccination by either direct plaque assay, or by virus amplification in Vero cells followed by plaque assay. For virus amplification using co-cultivation, 20 μL of serum in 2 mL of complete medium was used to infect Vero cells in 75 cm^2^ flask for 1h, then 20 mL medium was added, and incubation was continued for 48 h. Supernatant was harvested, and the virus was assayed by plaque assay.

In another set of experiments, C57BL/6J mice (5-6 week-old, Jackson Laboratories) were vaccinated with test vaccines (10^5^ PFU/mouse, s.c. into footpad) or PBS as described above.

Challenge was performed in the ABSL3 facility by inoculating C57BL/6J mice with a wild type CHIKV strain SL15649 into the footpad (s.c., 10^6^ PFU) at 49 days post vaccination. Animals were monitored daily for clinical signs including foot swelling and body weigh changes for 21 days post challenge.

### Western blot and IFA

CHIKV-specific antibody response was determined using sera collected on day 28 post vaccination by Western blot and indirect immunofluorescence assay (IFA). Mouse serum was probed in western blot with the lysates of CHIKV 181/25 infected Vero cells at 1:100 dilution, followed by alkaline phosphatase (AP)-conjugated goat anti-mouse IgG (H+L) secondary antibody (1:1000 dilution). For IFA, Vero cells seeded in chamber slides were infected with V5040 at MOI 0.01. At 24 h post-infection, Vero monolayers developed foci of V5040-infected cells. Monolayers were fixed with acetone and probed with mouse serum diluted 1:25, followed by Fluorescein isothiocyanate (FITC)-conjugated goat anti-mouse IgG (H+L) secondary antibody (1:25 dilution). Nuclei were stained with VectaShied mounting medium containing propidium iodide (Vector Laboratories, Inc., Newark, CA).

### Plaque reduction neutralization test (PRNT_50_)

Neutralizing antibody was determined using PRNT_50_, as well as by using nLuc-expressing CHIKV. Briefly, C57BL/6J mice (n=8-10/group equal number of sex) were sacrificed at various time points (3, 7, 14, 21, and 41 day post-vaccination) and the total blood was collected by terminal bleeding. For PRNT, diluted homologous virus was mixed with the same volume of serially diluted sera (1: 64 to 1: 2048, two-fold) and incubated at 37 °C for one hour. Virus-serum mixture was added to Vero 76 cells confluently grown in 24-well plates and incubated at 37 °C for 1h for virus adsorption. Cells were washed with PBS once, and overlayed with an overlay medium (1X EMEM with 2% FBS, 0.75% methylcellulose). After four days of incubation in a CO_2_ incubator at 37 °C, cells were fixed and viral foci were visualized with a crystal violet staining solution (2% paraformaldehyde, 10% ethanol, and 1% crystal violet). Number of viral foci forming units were normalized based on the average from the mock group and PRNT_50_ was calculated with a dose - response analysis with a four-parameter logistic model (XLfit, IBDS, UK). Statistical analysis was performed with GraphPad Prism (9.4.1).

Additionally, neutralization test was performed by using CHIKV expressing nLuc gene. Test sera were heat-treated at 56 °C for one hour then serially diluted in PBS at a starting dilution of 1:32. Thirty microliter of diluted serum was mixed with the same volume of diluted CHIKV-nLuc virus (600 pfu/30 μL, quadruplicates per sample). After an hour of incubation at 37 °C, 50 μL of the virus-serum mixture was added to Vero 76 cells grown overnight in a 96-well cell culture plate then further incubated in a 37 °C CO_2_ incubator for one hour for absorption. Then cells were washed with PBC once and replenished with the cell culture media (E-MEM with 10% FBS). Cells were incubated at 37 °C for two days and the nLuc activity from the virus replication was measured by using Nano-Glo® Luciferase Assay System (Promega) following the manufacturer’s protocol using a plate reader (HT4 Biotek). Luminescence data was normalized with values from wells with a normal mouse sera as 0% and mock infected control as 100%. PRNT_50_ was defined as the highest reciprocal dilution of the serial dilution series that produces > 50% reduction in the nano-luciferase activity.

## RESULTS

### Design of V5040 live-attenuated CHIKV with rearranged structural genes and 181/25 E2 mutations

The genetic structures of the CHIKV pMG5040 and V5040 RNA are schematically shown in Figure 1a. The full-length, functional RNA genome of the V5040 vaccine virus was encoded in the pMG5040 plasmid downstream from the optimized CMV promoter. The pMG5040 was prepared by re-designing the prototype CHIKV iDNA clone described elsewhere [25]. The pMG5040 plasmid is intended to launch replication of live-attenuated V5040 virus with rearranged genome in mammalian cells. After transfection of Vero cells with the pMG5040, the CMV promoter directs transcription of the functional viral genomic RNA starting from the 5’ terminus to the 3’-terminal poly-A sequence. Compared to the wild type CHIKV genome, the structural genes of V5040 RNA genome were rearranged. The CHIKV structural gene region was split into two open reading frames (ORFs), one expressing GP genes E3-E2-6K-E1, and the other expressing the C gene only. Each structural ORF was expressed from its own 26S subgenomic promoter, and included translational start and stop codons (Figure 1). Artificially rearranged genomes lead to the attenuation of many viruses and are resistant to reversions because many independent mutations would be required to restore the wild type virus sequence [32-34]. To further secure attenuation and safety of the experimental V5040 vaccine, mutations E2 Thr12Ile and Gly82Arg associated with attenuation of the CHIKV 181/25 vaccine were introduced in the GP region of the pMG5040 and V5040.

### Rearranged CHIKV V5040 replicates in Vero cells transfected with pMG5040 iDNA plasmid

Potentially, rearrangement of the genes can severely affect, or even prevent, replication of the CHIKV. To evaluate if V5040 virus can replicate, pMG5040 plasmid encoding rearranged CHIKV RNA genome was transfected in cultured cells as described in Materials and Methods. The pMG5040 was isolated from *E. coli* (Figure 1b). To launch V5040 CHIKV, pMG5040 plasmid was transfected into Vero cells (ATCC CCL-81.5). Unlike infection with CHIKV, which replicates in the cytoplasm, the transfection of pMG5040 leads to transcription of rearranged genomic RNA in the nuclei of transfected cells, and genomic RNA is transported to the cytoplasm where RNA translation and virus synthesis take place. After transfection, expression of replication competent virus was confirmed by plaque assay in the growth medium harvested from transfected cells. As seen in Figure 2, incubation of cells with the medium from pMG5040-transfected cells (see Materials and Methods) generated plaques under agarose overlay at 24-48 h post infection (Figure 2). Compared to control CHIKV 181/25-generated plaques, V5040 plaques were smaller in diameter, potentially reflecting additional attenuation of V5040 in comparison with 181/25 virus. The V5040 virus harvested from the supernatant at 48 h post-transfection had an infectious titer 10^8^ PFU/mL indicating successful rescue of replication-competent virus from pMG5040 iDNA plasmid and efficient replication of V5040 in Vero cells. Expression of CHIKV-specific antigens in V5040-infected cells was confirmed by IFA using CHIKV-specific antiserum VR-1241AF as described elsewhere [25].

**Figure 2.**
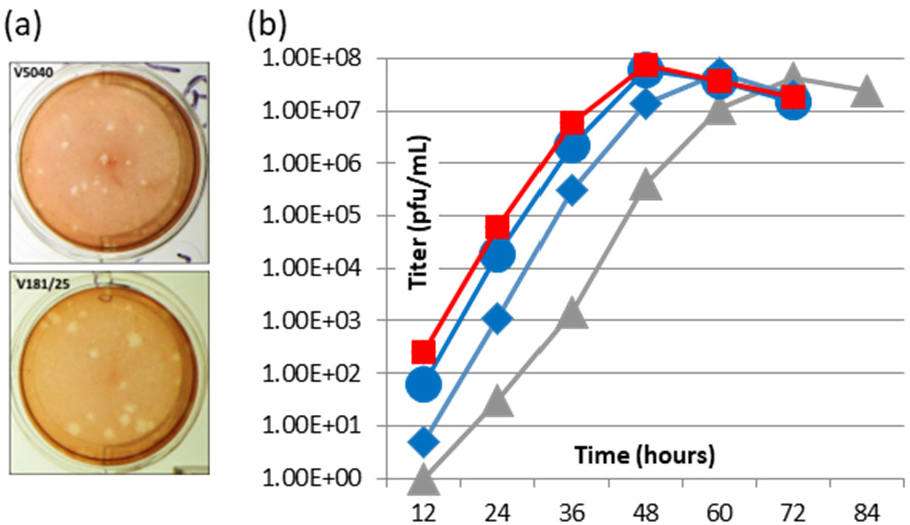
Preparation of V5040 CHIKV in Vero cells. (a) Plaque morphology of attenuated V5040 and 181/25 vaccine viruses. (b). Comparison of the growth kinetics of V5040 in Vero cell infected with V5040 (10^2^ PFU, blue diamonds; 10^3^ PFU, blue circles) or transfected with pMG5040 (100 ng, grey triangles). As control, CHIKV 181/25 vaccine virus was used (10^3^ PFU, red squares).

In the next experiment, we compared replication of V5040 virus in Vero cells infected either with the V5040 virus (100 PFU or 1000 PFU representing MOI of 10^-5^ and 10^-4^, respectively), or transfected with pMG5040 plasmid. As a control, we used experimental CHIKV 181/25 vaccine. Samples of the culture medium from transfected and infected cells were collected every 12 h and the viral titers were quantitated by plaque assay. As shown in Figure 2(b), infection at MOI of 10^-5^ resulted in effective virus replication comparable with CHIKV 181/25 kinetics and reached 10^8^ PFU/ml at 48 h post infection. Infection at lower MOI 10^-4^ had similar kinetics peaking at 60 h. p. i. In pMG5040-transfected cells, most cells expressed CHIKV antigens at 48 h post-transfection and infectious virus in culture medium approached 10^8^ PFU/ml at 72 h. The observed delay of the peak titer in pMG5040 transfected cells as compared to V5040 infected cells can be explained by the timing associated with cell recovery after pMG5040 electroporation and by delay associated with DNA penetrating the nuclei of transfected cells, transcription of RNA and transport into cytoplasm to start replication of V5040. Thus, replication of V5040 virus from plasmid is expected to be delayed causing peak titer at a later point as compared to the control CHIKV 181/25 virus. Similar observation was made with DNA-launched Venezuelan equine encephalitis virus (VEEV), a related rearranged alphavirus [35]. Based on our experiments, we observed that infection 100-1000 PFU of V5040 virus, or transfection with 100 ng of pMG5040 iDNA efficiently initiated replication of live attenuated CHIKV *in vitro*.

### Rearranged V5040 virus is immunogenic in mice

To evaluate safety and immunogenicity of rearranged V5040 CHIKV vaccine, we used BALB/c and C57BL/6 mice. In the first experiment, BALB/c mice were vaccinated with a single s.c. dose of 10^4^ PFU or 10^5^ of V5040 virus prepared from the growth medium of pMG5040-transfected Vero cells. Similarly, 181/25 virus was administered as control. After injections, all mice remained healthy with no detectable adverse effects such as changes in weight or behavior. Serum samples were collected as described in Materials and Methods. Viremia was not detectable in V5040-vaccinated mice by direct plaque assay (detection limit 25 PFU/ml). However, in 40% of the vaccinated mice, low viremia was detected by co-cultivation assay (Table 2). At day 28, mice seroconverted as determined by IFA and western blot (Table 2; Figure 3). Similarly, BALB/c mice vaccinated with a single s.c. dose of 10^5^ PFU of V5040 virus did not show any safety concerns and developed antibody response (Table 2).

**Table 2.** Immunogenicity of V5040 CHIKV vaccine in BALB/c and C57BL/6 mice.

|  | Mouse Model | Dose, PFU (route) | Viremia |  | Serum Antibody |
| --- | --- | --- | --- | --- | --- |
|  |  |  | Direct | Amplification |  |
| Exp. 1 | BALB/c | $10^4$ (s.c.) | - | + | + (WB, IFA, PRNT<sub>50</sub>) |
| | | $10^5$ (s.c.) | | | |
| Exp. 2 | C57BL/6J | $10^5$ (s.c.) | nt | nt | + (ELISA, PRNT<sub>50</sub>) |

PFU, plaque-forming units; s.c., subcutaneously; WB, western blot; IFA, indirect immunofluorescence assay; PRNT<sub>50</sub>, plaque reduction neutralization assay; nt, not tested.

**Figure 3.**
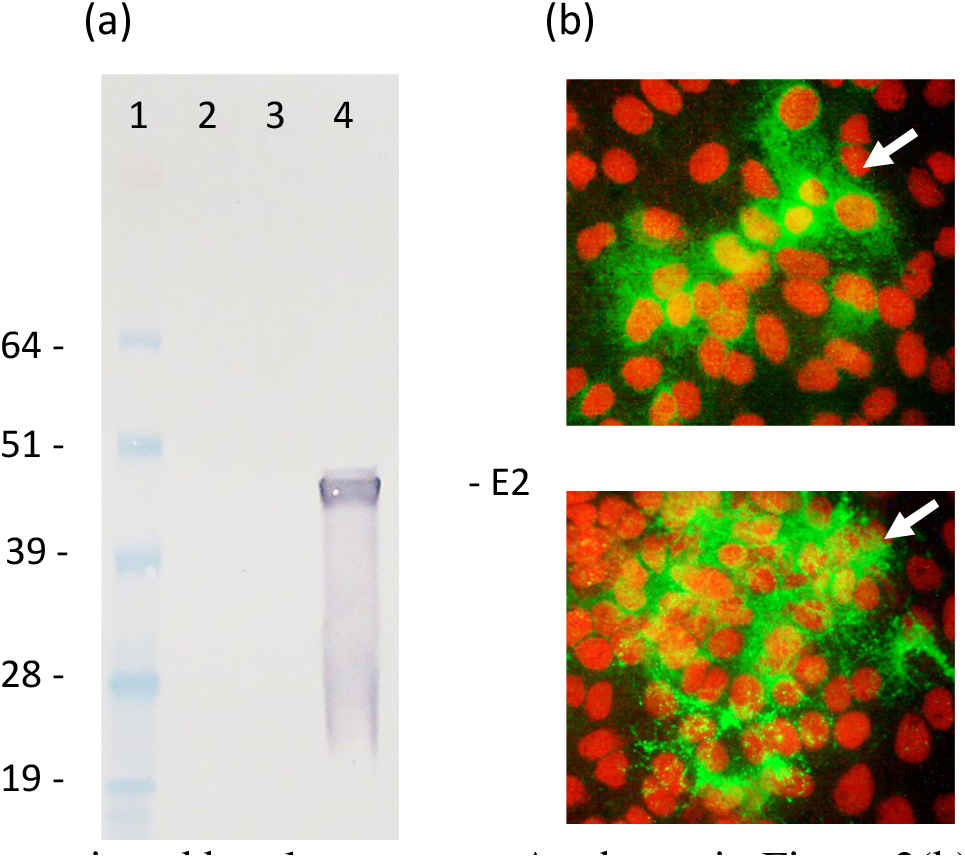
Immunogenicity of V5040 virus in mice. (a) western blot and (b) IFA with antisera from BALB/c mice vaccinated with V5040 virus. (a) For western blot, mouse serum was probed at 1:100 dilution followed by AP-conjugated goat anti-mouse IgG (H+L) secondary antibody (1:1000 dilution). Lane 1, SeeBlue Plus2 standard, Lane 2, control Vero cell lysate; Lane 3, blank; Lane 4, lysate of Vero cells infected with CHIKV 181/25 virus. Predicted band of E2 is indicated. (b) For IFA, Vero cells seeded in chamber slides were infected with V5040 at MOI 0.01. At 24 h post-infection, Infected monolayer cells were fixed with acetone and probed with mouse serum diluted 1:25, followed by FITC-conjugated goat anti-mouse IgG (H+L) secondary antibody (1:25 dilution). Infected Vero monolayers developed foci of V5040-infected cells shown in green (indicated with arrows). Nuclei are stained in red using VectaShield mounting medium containing propidium iodide.

Next, we evaluated neutralization activity of V5040 vaccine with antisera from vaccinated C57BL/6 mice using a standard PRNT_50_ assay, as well as neutralization of CHIKV-nLuc virus. As control, we used 181/25 vaccine. In a standard PRNT_50_ assay, both vaccines demonstrated measurable neutralization activity starting from 7 days post vaccination after a single dose of vaccine (Figure 4). Compared to 181/25 CHIKV vaccine, iDNA-based V5040 vaccine induced stronger and more durable PRNT_50_ neutralizing titers. While approximately 50% (4-5 / 8 mice, per group) of 181/25 vaccinated mice showed a measurable neutralization activity (>1:32), nearly all of V5040 vaccinated mice demonstrated PRNT_50_ > 1:128 after 14 days post vaccination, with maximum titer of 1:512. Interestingly enough, V5040 showed a better longevity of neutralization activity, showing a significantly higher PRNT_50_ titer compared to CHIKV 181/25 at 42 days post vaccination (p < 0.0007, Two-way ANOVA, mixed effect model).

**Figure 4.**
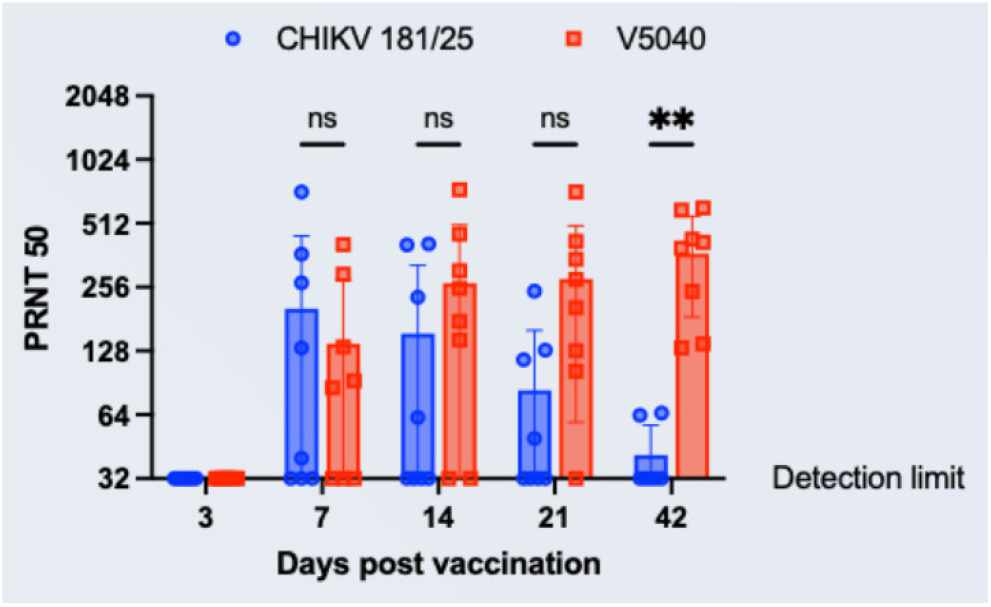
Neutralization antibodies from C57BL/6 mice vaccinated with V5040 or CHIKV 181/25. Antisera collected from terminal bleeding of mice vaccinated either with V5040 or CHIKV 181/25 were tested for PRNT_50_ assays against their corresponding homologous virus. Blue circles and red squares indicate antisera of CHIKV 181/25 and V5040, and each symbol represents serum from an individual mouse (n=8/group/timepoint). The bar and whiskers indicate the means and ±their standard deviation. Ns: not significant. **, P < 0.0021, Two-Way ANOVA with Sidak’s multiple comparisons test (Prism).

CHIKV-neutralizing effect of serum from vaccinated mice was confirmed in a CHIKV-nLuc virus-based neutralization experiment (Figure 5). Neutralization was detected in both V5040 and 181/25 vaccinated sera, with highest titers up to 1:1024 in V5040 vaccinated mice. Again, the V5040-vaccinated mice appeared to have higher titer of neutralization, with p=0.0152 between V5040 and 181/25 groups (Figure 5). The observed titers are expected to be protective, as PRNT_50_ ≥10 may be sufficient to correlate with protection from symptomatic infection and subclinical seroconversion [36]. Challenge was performed by inoculating a wild type CHIKV strain SL15649 into the footpad s.c., 10^6^ PFU at 49 days post vaccination. Foot swelling and protection after challenge were detected; however, it was statistically not significant due to the limitation in the animal model (i.e., limited foot swelling in adult mice, > 4-6 weeks old).

**Figure 5.**
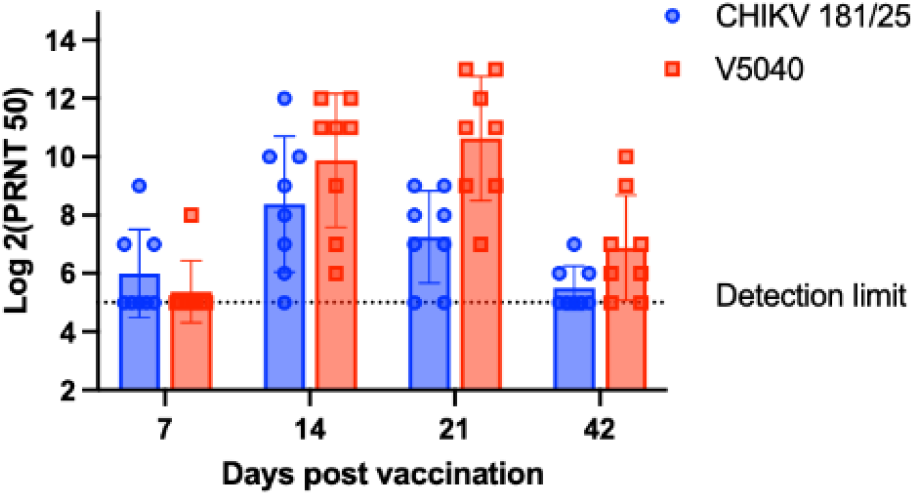
Anti-CHIKV neutralization activity of antisera immunized with CHIKV 181/25 and V5040 vaccine. Diluted serum was mixed with the same volume of diluted CHIKV-nLuc virus (600 pfu) for 1h, then used to infect Vero 76 cells in a 96-well cell culture plate for two days. The nLuc activity from the virus replication was measured by using Nano-Glo® Luciferase Assay System (Promega) following the manufacturer’s protocol using a plate reader (HT4 Biotek). Each data points represents a group of 8 mice (4 male and 4 female), with p=0.0152 between 181/25 and V5040 groups. The statistical analysis was performed with GraphPad using two-way ANOVA.

In summary, although additional experiments are needed to evaluate protection and potential difference in magnitude of the neutralizing response between two highly attenuated viruses, the results clearly indicate high level of neutralization activity of serum from mice vaccinated with rearranged V5040 CHIKV.

## DISCUSSION

Live attenuated vaccines represent approximately half of all licensed vaccines in the U.S. [37]. Live vaccines Zostavax, RotaTeq (Merck), FluMist (AstraZeneca), Rotarix (GSK) have been recently approved showing that live attenuated platform can be configured to meet stringent FDA safety standards.

Because of its clinical history, the 181/25 vaccine is a good starting point for CHIKV vaccine development. Previously, we developed iDNA approach as a novel infectious clone plasmid, in which the full-length viral RNA genome is transcribed from the CMV promoter [24, 30, 38]. Using iDNA approach, we prepared the iDNA encoding the prototype 181/25 CHIKV vaccine [25]. The latter encoded 181/25 genomic RNA in the plasmid downstream from CMV promoter. This plasmid launched the CHIKV 181/25-like vaccine virus in cell culture and *in vivo*. Vaccination of BALB/C mice with this 181/25 iDNA plasmid protected mice from CHIKV challenge [25]. The attenuating mutations in the iDNA-generated virus were confirmed by DNA sequencing, with no reversions detected [39]. Additionally, the iDNA-derived virus is expected to have lower heterogeneity (% SNPs) at the attenuating-mutation sites, E2-12 and E2-82, as compared to the prototype 181/25 virus. In a previous study, the iDNA-derived RNA had fewer SNPs vs. the 181/25 virus vaccine, suggesting potential safety advantage vs. classic live-attenuated virus [39].

Genetic rearrangements potentially can be utilized to engineer safer live-attenuated vaccines [26]. Gene rearrangement is expected to be highly resistant to reversion because multiple independent mutations would be needed to restore wild-type genotype. Here, we applied genetic rearrangement for the development of novel CHIKV live-attenuated vaccine based on an investigational 181/25 live-attenuated CHIKV vaccine. In early studies, experimental CHIKV vaccine clone 181/25 was reported as highly immunogenic in Phase II clinical trial; however, mild transient arthralgia was observed in some patients [23]. Vaccine was evaluated in 59 healthy volunteers, with 98% seroconversion rate. However, up to 8% experienced mild transient arthralgia [23]. Reversion mutations have been detected in viremic patients that received the 181/25 IND vaccine [40] indicating the need for safety improvement. The structural gene rearrangement described in this study represents a mutation that is resistant to reversion.

In the pMG5040 iDNA plasmid, the DNA copy of the genome of CHIKV vaccine virus is rearranged, with the capsid gene placed downstream from the glycoprotein genes using a duplicate subgenomic promoter. This gene rearrangement does not affect the protein sequence and the immunogenic epitopes. However, until now it was not known if rearrangement can be engineered for CHIKV 181/25 live-attenuated vaccine without impairing its fitness in vitro and in vivo. Therefore, in this study, we showed that V5040 virus efficiently replicates in pMG5040-transfected Vero cells. Finally, safety and immunogenicity was shown by vaccinating mice with V5040.

In another study, we also prepared a rearranged V4020 vaccine from the alphavirus vaccine TC83 for Venezuelan equine encephalitis virus (VEEV) [35]. V4020 vaccine was safe, immunogenic, and efficacious in mice and non-human primates [35, 41, 42] suggesting genetic rearrangement can be used for distinct alphavirus vaccines to improve safety.

Taken together, our results showed the feasibility of iDNA approach to prepare CHIKV and other alphavirus live vaccines with rearranged genome for higher safety. Potentially, iDNA vaccine can be used for vaccination directly, by injection into muscle of the vaccine recipient. Such iDNA vaccine would combine the benefits of conventional DNA immunization except it uses small quantities of DNA to launch efficacious live-attenuated vaccines [24]. The iDNA vaccine turns a small number of cells in muscle of vaccine recipients into cell-scale vaccine “factories”. The iDNA uses a well-established manufacturing technology of bacterial production of plasmids, which are easier to bank, control, and manipulate vs. live virus stocks. Since iDNA plasmids represent genetically defined homogenous clones that after vaccination make the virus that undergoes minimal replication cycles, the probability of reversion mutations compared to traditional manufacturing is reduced [24]. The iDNA vaccination may also have additional advantage for immunogenicity due to immunostimulatory effects of DNA vaccine. In addition to improved safety and immunogenicity, a CHIKV iDNA vaccine offers the potential advantages of high purity, genetic stability, simplicity of production, no cold chain, single-dose vaccination, and long-lasting immunity [24]. Thus, iDNA would combine the advantages of DNA immunization and the high efficacy of live-attenuated vaccines.

Additional research is needed on V5040 or pMG5040 as CHIKV vaccines. In the proof-of-concept experiments, we showed that vaccination of BALB/C mice with the iDNA-based 181/25 vaccine protected them from CHIKV challenge [25]. However, pMG5040 has not yet been tested in an iDNA format for vaccination. In addition, protective effect of V5040 vaccine needs to be tested. Potentially, neurovirulent CHIKV Ross strain can be used for this purpose [25], as well as suckling mice and mice with interferon-defective genotype [43]. Furthermore, advanced safety, immunogenicity, and efficacy studies in non-human primates will be also needed for pre-clinical testing of CHIKV vaccine. Non-human primates have been successfully used to evaluate the safety and immunogenicity of live CHIKV and other vaccines [44]. Another future project is to confirm that no reversion mutations occur *in vivo*. To address this, genetic stability studies *in vivo* (multiple passages in tissues and adult mouse brains) can be carried out to provide additional safety information as previously described [42].

Several other experimental CHIKV vaccines have been described. These include live chimeric alphaviruses that carry CHIKV structural proteins [45], as well as formalin-inactivated vaccine adjuvanted with aluminum hydroxide [46]. Furthermore, a virus-like particle (VLP) CHIKV vaccine protected nonhuman primates from CHIKV challenge [47, 48] and is currently in the clinical trials. VLP vaccine has achieved immunogenicity milestones in the clinical trial. DNA-based vaccines are another promising vaccination strategy [49]. However, many of experimental vaccines required two or more vaccinations to elicit protective immune response, which could be a disadvantage when rapid control of an outbreak is needed. A single-dose CHIKV vaccine that protects for the long-term would be a major benefit to global health. Measles virus-vectored CHIKV vaccine is currently in the clinical trials [50]. Also, successful completion of the lot-to-lot Phase 3 trial of single-dose chikungunya vaccine candidate was announced, CHIKV Δ5nsP3 (VLA1553) [51].

The proposed rearranged iDNA-derived V5040 vaccine is designed to improve safety and induce effective immunity after a single injection. Live-attenuated vaccine has a favorable cost/benefit ratio and includes innovative safety features. It can be used to vaccinate individuals at risk of CHIKV infection, as well as for rapid deployment in outbreak situations to immunize against CHIKV in endemic and non-endemic areas.

## ACKNOWLEDGEMENTS

Authors sincerely thank Elena Klyushnenkova, Alex Tibbens and Noble Life for experimental contributions. Research reported in this publication was supported by the National Institute of Allergy and Infectious Diseases of the National Institutes of Health under Award Number R43AI152717. The content is solely the responsibility of the authors and does not necessarily represent the official views of the National Institutes of Health.

